# High-resolution architecture of human epiphysis formation

**DOI:** 10.1101/2020.10.13.337733

**Authors:** Heng Sun, Ya Wen, Weiliang Wu, Tian Qin, Chengrui An, Chunmei Fan, Yishan Chen, Junfeng Ji, Ting Gang Chew, Jiansong Chen, Hongwei Ouyang

## Abstract

Human limb skeletal system consists of both bone and cartilage which originated from fetal cartilage. However, the roadmap of chondrocyte divergent differentiation to bone and articular cartilage has yet to be established. Epiphysis possesses articular cartilage, growth plate and the secondary ossification center (SOC), making it an ideal model to uncover the trajectory of chondrocyte divergent differentiation. Here, we mapped differentiation trajectory of human chondrocyte during postnatal finger epiphysis development by using single-cell RNA sequencing. Our results uncovered that chondroprogenitors have two differentiation pathways to hypertrophic chondrocytes during ossification, and one pathway to articular chondrocytes for formation of cartilages. Interestingly, we found that, as an addition to the known typical endochondral ossification path from resting, proliferative to hypertrophic chondrocytes, there was a bypass by which chondroprogenitors differentiate into hypertrophic chondrocytes without proliferative stage. Furthermore, our results revealed two new chondrocyte subpopulations (bypass chondrocytes as it appeared in the ossification bypass, and *ID1*^+^ chondroblasts in articular chondrocyte path) during postnatal epiphysis development in addition to six well-known subpopulations. Overall, our study provides a comprehensive roadmap of chondrocyte differentiation in human epiphysis thereby expanding the knowledge of bone and articular cartilage, which could be utilized to design biotherapeutics for bone and articular cartilage regeneration.

## Human chondrocyte identification

To investigate the chondrocyte differentiation trajectory during human epiphysis development, the phalanges from polydactyl patients were used. The histological staining of the 2-year-old polydactyl phalange showed a typical long bone structure (Fig. S1A), similar with the human femur(*1*).

The bone and cartilage tissues from polydactyly samples were collected and digested for 4 hours before single-cell isolation, library preparation and sequencing (Fig. 1A). After quality control process, we obtained 27,461 single cells for data analysis (Fig. S1B). We clustered these cells using the well-established method Seurat(*2*, *3*), and found 10 clusters identified as chondrocytes (*COL2A1*),fibroblasts (*COL1A2*), vascular cells (*PECAM1*), muscular cells (*ACTA2*), antigen presenting cells (*HLA-DRA*), osteoblasts (*BGLAP*), natural killer cells (*NKG7*), Schwann cells (*MPZ*), erythrocytes (*HBA1*), and megakaryocytes (*MMRN1)* (Fig. S1C and S1D). Among them, both clusters 0 and 5 robustly expressed the chondrocyte marker *COL2A1,* and they were highly related with cartilage development (Fig S1E). Therefore, we selected these 14,434 single cells from clusters 0 and 5 for further analysis.

**Figure 1:**
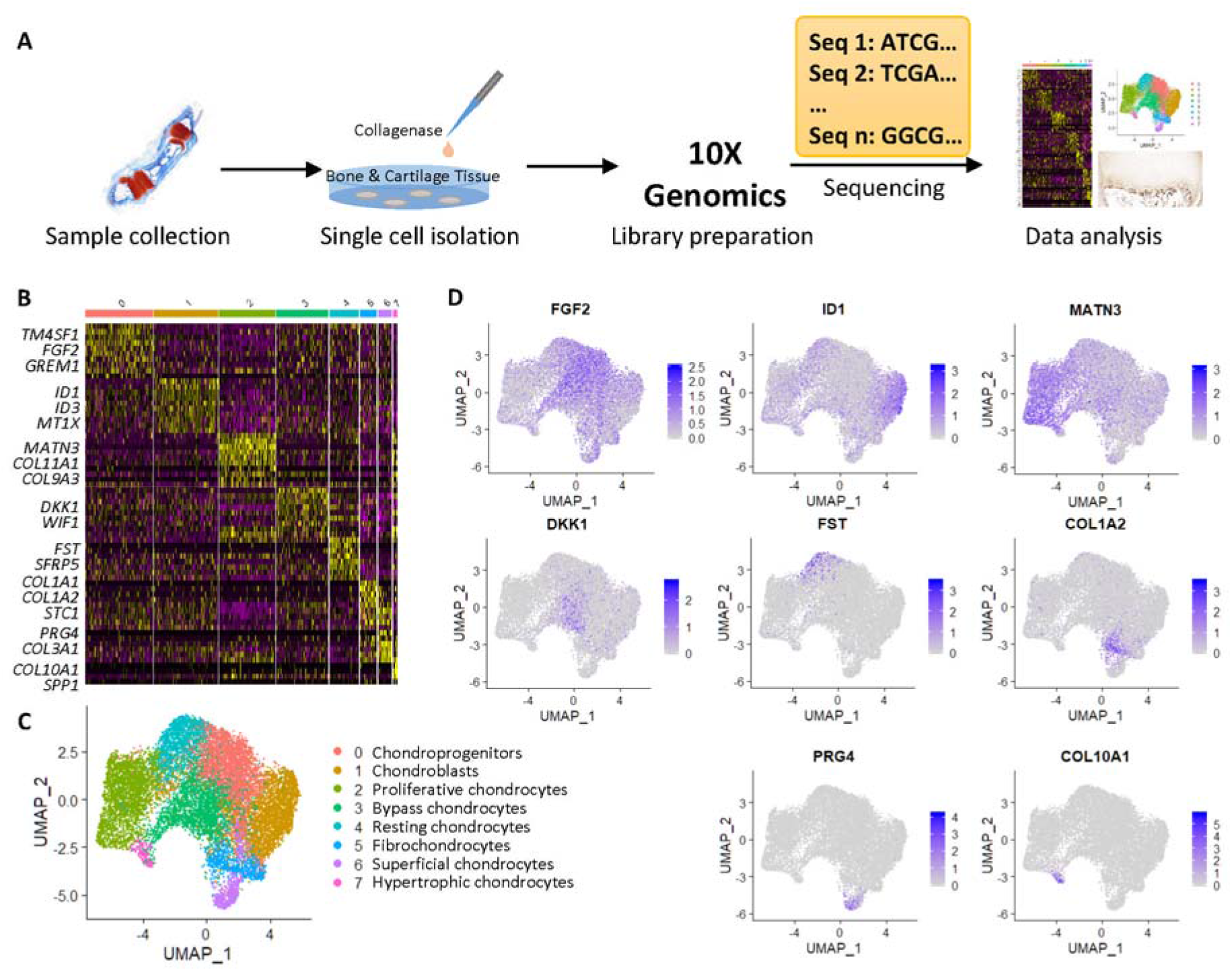
Chondrocyte heterogeneity. (**A**) Workflow of the single-cell RNA sequencing analysis. (**B**) Heatmap of the differentially expressed genes for each cluster. (**C**) Cell clusters visualized by UMAP. (**D**) Expression levels of the representative genes for each cluster.

Using Seurat, the cartilage-related cells were clustered into eight clusters (Fig. 1B and C). With the highly expressed genes in each cluster, we were able to identify the well-known chondrocyte subpopulations, including chondroprogenitors (*FGF2*), resting chondrocytes (*FST*), proliferative chondrocytes (*MATN3*) hypertrophic chondrocytes (*COL10A1*), fibrochondrocytes (*COL1A2*), and superficial chondrocytes (*PRG4).* Interestingly, we also discovered new transitional cells, like the *DKK1^+^* cluster, and *ID1*^+^ cluster which would be discussed later (Fig. 1C and D).

## Endochondral ossification fate of human chondrocytes

In an attempt to map the trajectory of chondrocyte differentiation, we performed the pseudo-temporal analysis by using Monocle3 (*4–6).* Our analysis results showed that chondrocyte differentiation started from chondroprogenitors (dark purple) and eventually adopted either articular or hypertrophic cell fate (yellow) by progressing through different paths on the pseudo-time axis (Fig. S2A).

We first looked into the endochondral ossification branch, in which chondroprogenitors differentiate into resting chondrocytes, proliferative chondrocytes and eventually hypertrophic chondrocytes in a step-wise manner (Fig. 2A and B). The chondroprogenitors highly expressed *FGF2* encoding the growth factor which is important for cell proliferation and tissue development. *FGF2* is expressed in human fetal cartilage and its expression is located at resting and proliferative zones, but not hypertrophic zone(*7*). *TM4SF1* and *GREM1,* the two mesenchymal stem cell markers(*8*, *9*), were also highly expressed in this cluster, confirming the progenitor identity (Fig. 2C and S2B). Immunostaining of the 2-year-old middle phalange proximal epiphysis showed that, in line with *FGF2,* the Transmembrane 4 L6 Family Member 1 (encoded by *TM4SF1)* was also abundantly expressed in resting and proliferative zones but decreased in hypertrophic zone (Fig. 2C).

**Figure 2.**
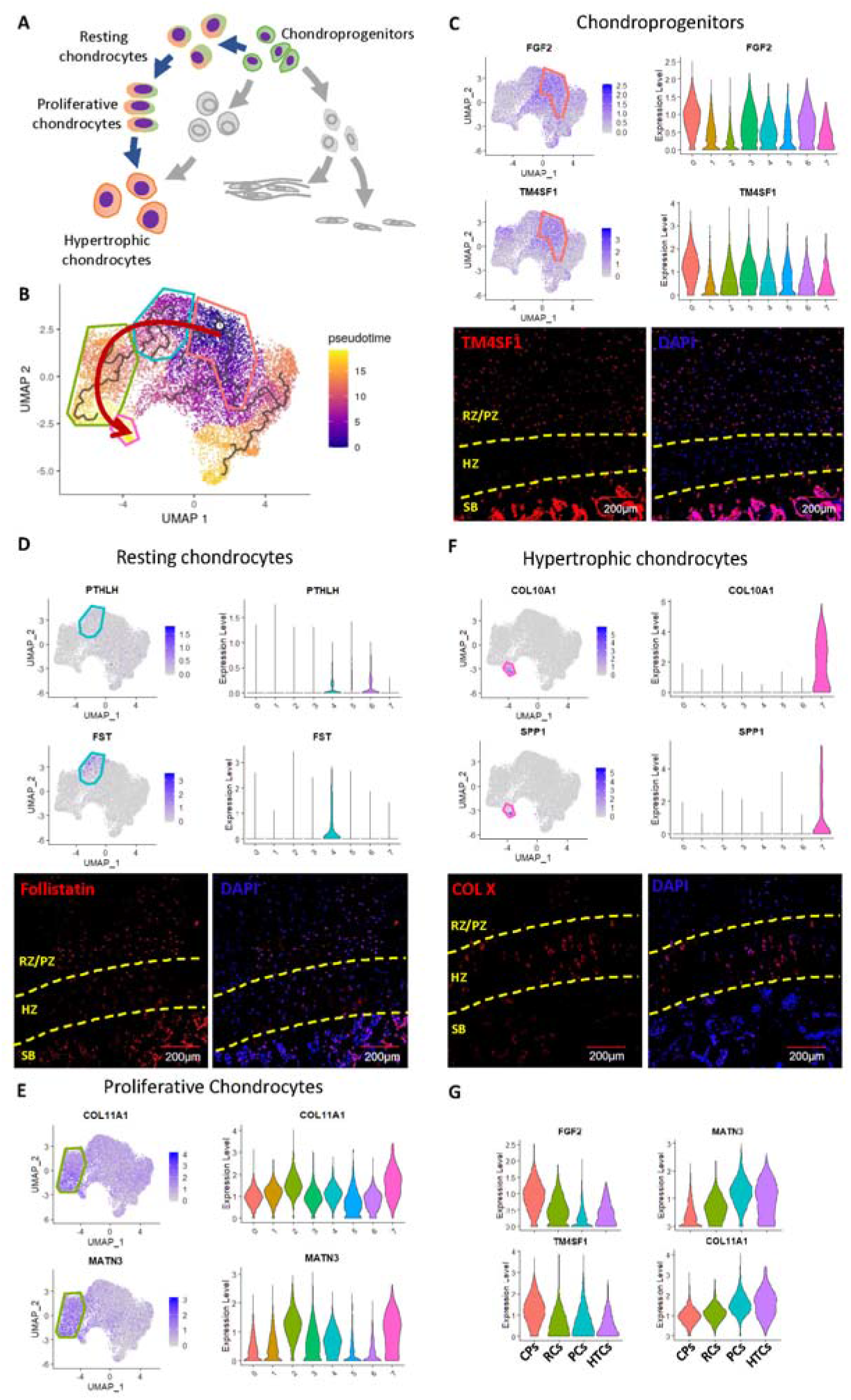
Identification of the typical endochondral ossification trajectory. (**A**) Schematic of the typical endochondral ossification trajectory. (**B**) Pseudotime analysis visualized by UMAP. Red arrow shows the typical endochondral ossification trajectory. (**C-F**) Gene expression levels and immunostaining of the chondroprogenitor (C), resting chondrocyte (D), proliferative chondrocyte (E), and hypertrophic chondrocyte (F) markers. RZ/PZ, resting zone/proliferative zone; HZ, hypertrophic zone; SB, subchondral bone. (**G**) Gene expression trends in the typical endochondral ossification trajectory. CPs, chondroprogenitors; RCs, resting chondrocytes; PCs, proliferative chondrocytes; HTCs, hypertrophic chondrocytes.

Resting chondrocytes were marked by *PTHLH,* the gene that encodes parathyroid hormone-related protein (PTHrP), and *SFRP5,* which were both reported to be expressed in resting chondrocytes(*10*, *11*). Interestingly, we found that this chondrocyte subpopulation expressed another marker gene *FST* which encodes follistatin (Fig. 2D and S2C), a BMP antagonist. Immunostaining of the phalangeal epiphysis demonstrated that follistatin positive resting chondrocytes were located near the proliferative and hypertrophic zones (Fig. 2D). These results are consistent with the previously reported inhibition of BMP signaling in resting zone(*12*).

Proliferative chondrocytes and hypertrophic chondrocytes had the highest expression levels of the mature cartilage matrix genes *COL9A3, COL11A1* (*13*) as well as *MATN3* and *COL6A3* (Fig. 2E and S2D). The expression of the pre-hypertrophic marker *PTH1R* was elevated in the proliferative chondrocytes too, consistent with previous studies that the chondrocytes in the bottom of the proliferative zone would become hypertrophic (Fig. S2E). The hypertrophic markers *COL10A1, SPP1* and osteogenic marker *IBSP* were all highly expressed in the hypertrophic chondrocytes, demonstrating the terminal chondrocyte differentiation. As expected, the immunostaining of the phalangeal epiphysis showed the positive signals of type X collagen right next to bone (Fig. 2F and S2F).

The four chondrocyte subpopulations exhibited the typical endochondral ossification trajectory from chondroprogenitors to hypertrophic chondrocytes in the growth plate, which was consistent with the conventional view (*14*).The gene expression trend along this trajectory also confirmed that the expressions of progenitor-related genes *FGF2* and *TM4SF1* were declined gradually with concomitant step-wise upregulation of the mature chondrocyte-related extracellular matrix genes *MATN3* and *COL11A1* (Fig. 2G). Therefore, our data successfully recapitulated the endochondral ossification trajectory.

## Articular chondrocyte differentiation fate of human chondrocytes

The articular chondrocyte differentiation trajectory was composed of chondroprogenitors, the newly discovered *ID1^+^* chondroblasts, fibrochondrocytes and superficial chondrocytes (Fig. 3A and B).

**Figure 3.**
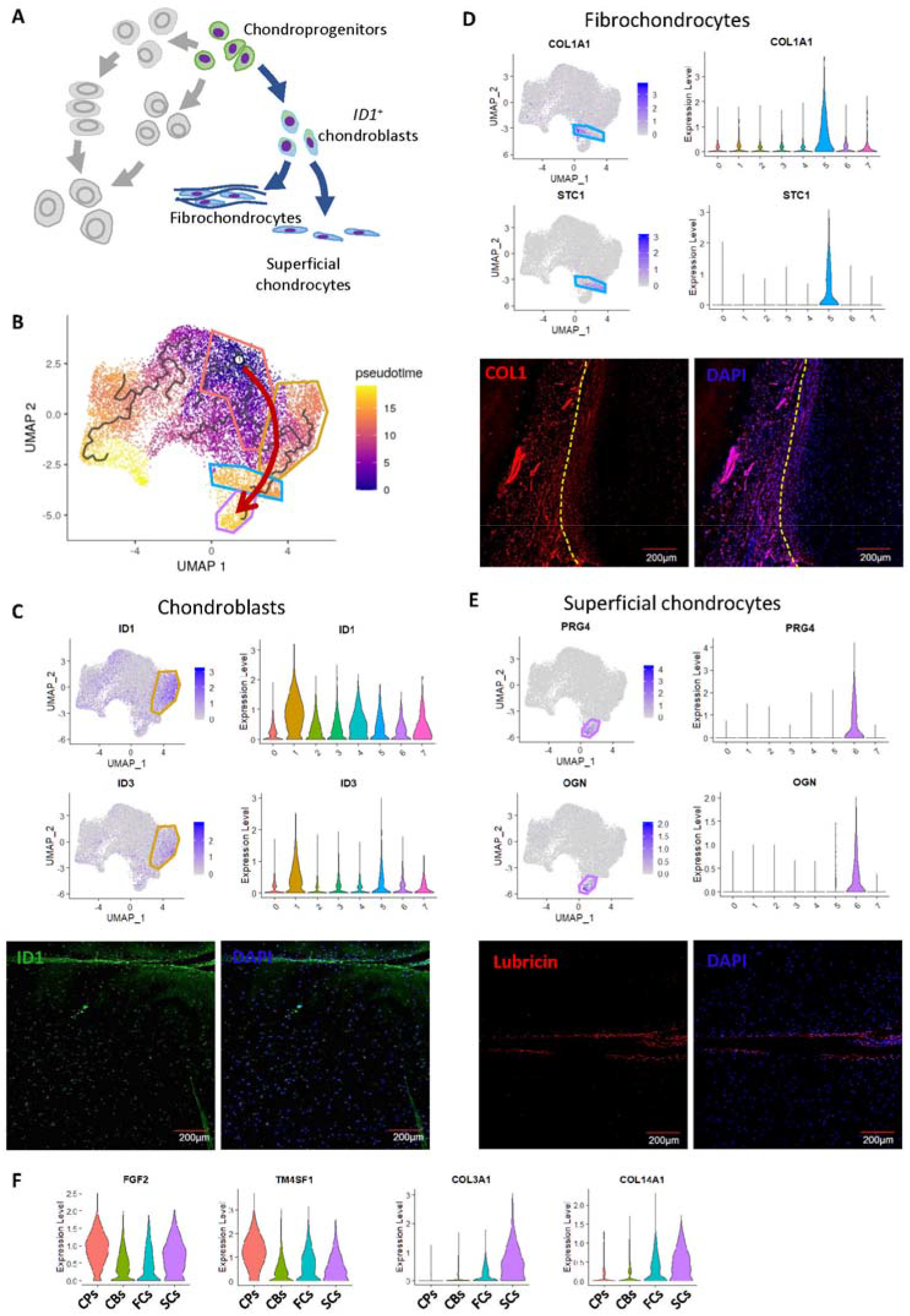
Identification of the articular chondrocyte differentiation trajectory. (**A**) Schematic of the articular chondrocyte differentiation trajectory. (**B**) Pseudotime analysis visualized by UMAP. Red arrow shows the articular chondrocyte differentiation trajectory. (**C-E**) Gene expression levels and immunostaining of the chondroblast, fibrochondrocyte, and superficial chondrocyte markers. (**F**) Gene expression trends in the articular chondrocyte differentiation trajectory. CPs, chondroprogenitors; CBs, chondroblasts; FCs, fibrochondrocytes; SCs, superficial chondrocytes.

*ID1* and *ID3,* the two markers of the chondroblast subpopulation (Fig. 3C and S3A), were reported to be expressed in less differentiated chondrocytes, and play a key role in the regulation of cell-cycle progression and cell differentiation in chondrocyte and other cells (*15*–*17*). *ID1* was also up-regulated in mesenchymal stem cells forming cartilage(*18*). The immunostaining of the phalangeal epiphysis showed a diffused distribution of ID1 in the periarticular area, indicating that these cells may give rise to the articular cartilage (Fig. 3C and S3A). As this subpopulation was located between chondroprogenitors and the fully differentiated chondrocytes, we regarded this subpopulation as chondroblasts (*19*).

*COL1A1/A2* marked the fibrochondrocytes (Fig. 3D and S3B). Since there are soft tissues including perichondrium, synovium and tendon that connect to the cartilage, it is not surprising to find the chondrocytes with soft connective tissue matrix in the transition section as shown by the immunostaining results. In fact, *STC1* and *COL14A1* were also found to be highly expressed in fibrochondrocytes (Fig. 3D and S3C). Given the known expressions of *STC1* and *COL14A1* in human synovium and tendon, respectively, (*20*, *21*) these data suggested that this subpopulation formed the cartilage that connected adjacent tissues.

The superficial chondrocytes were labeled by *PRG4,* the well-known marker for chondrocytes in the superficial layer. *Prg4^+^* cells were also thought to possess progenitor or regeneration potential(*22*–*24*). Consistently, immunostaining showed that the expression of lubricin encoded by *PRG4* was also located at the superficial layer (Fig. 3E and S3D), and superficial chondrocyte subpopulation had higher *FGF2* and *GREM1* levels compared with the adjacent chondroblasts and fibrochondrocytes (Fig. 2C and S2B). Moreover, *OGN,* the gene that encodes osteoglycin, was highly expressed in the superficial chondrocytes as well, which is in line with a previous report(*25*) (Fig. 3E).

Consistent with the endochondral ossification trajectory, the expression levels of the progenitor-related genes *FGF2* and *TM4SF1* decreased progressively, while the extracellular matrix-related genes *COL3A1* and *COL14A1* were increased (Fig. 3F).

Taken together, these data demonstrated the articular chondrocyte differentiation trajectory, improved the understanding of the articular cartilage development in the epiphysis.

## The bypass of ossification for human chondrocyte differentiation

Intriguingly, in the pseudotime trajectory we found a cell type that bypasses the conventional step-wise differentiation of human chondrocytes. It appears that the chondroprogenitors differentiated to hypertrophic chondrocytes directly after transiting into a specific chondrocyte subpopulation. Therefore, we named this subpopulation as bypass chondrocytes (Fig. 4A and B). Both *DKK1* and *WIF1* encoding antagonists of WNT signaling pathway were highly expressed in this subpopulation together with *SMPD3* (Fig. 4C and S4A). *SMPD3* is expressed in pre-hypertrophic chondrocytes, marking the hypertrophic fate of the cells(*26*). *Smpd3* deficiency caused ossification retardation and SOC absence(*27*), pointing to the significant role of *SMPD3* in the epiphysis development. Same as the previous trajectory, the chondrocytes lost the chondroprogenitor identity and become hypertrophic gradually, as *FGF2* and *TM4SF1* expression were down-regulated and *PTH1R* and *SMPD3* were up-regulated (Fig. 4D).

**Figure 4.**
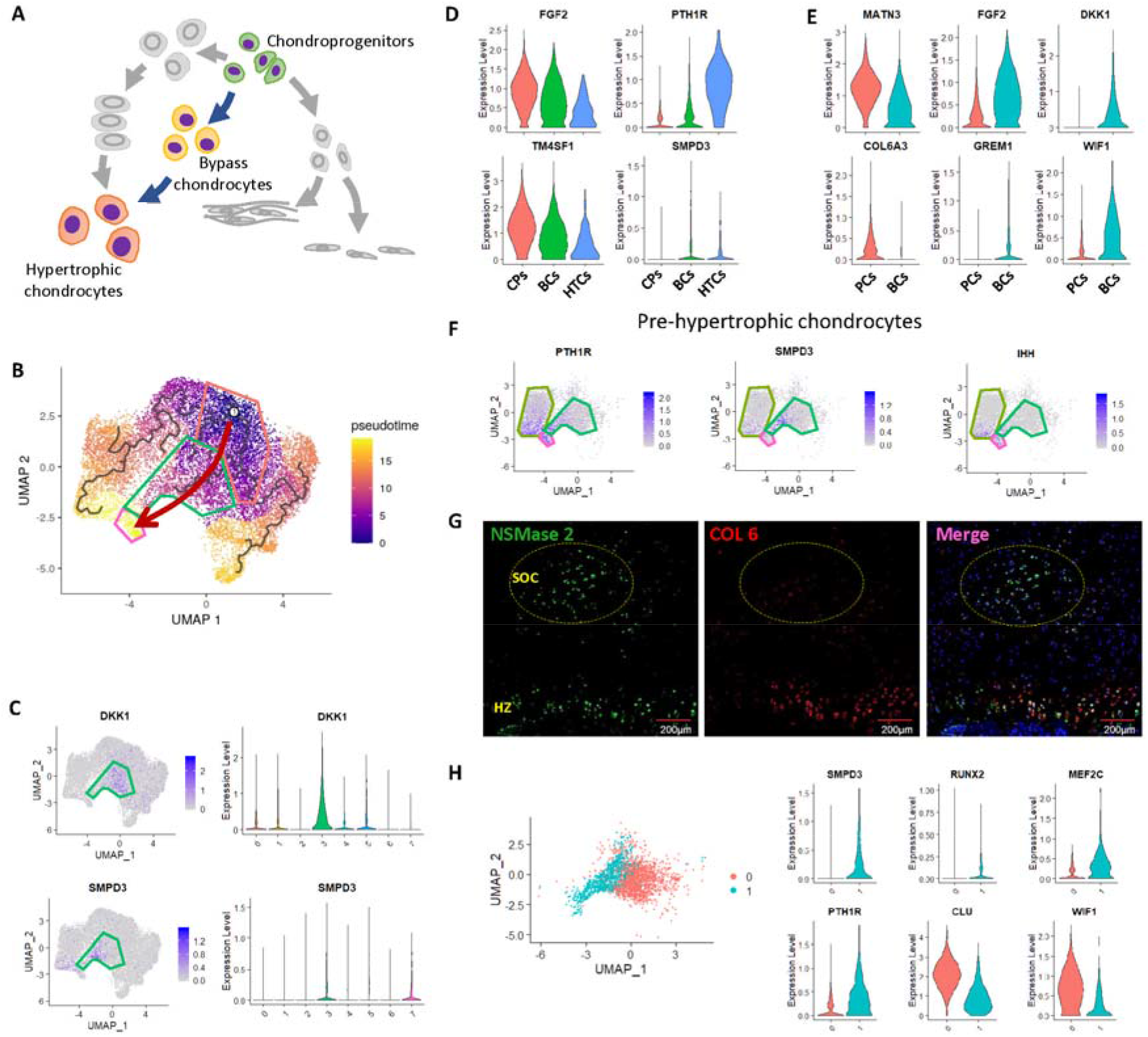
Identification of the chondrocyte differentiation bypass. (**A**) Schematic of the chondrocyte differentiation bypass. (**B**) Pseudotime analysis visualized by UMAP. Red arrow shows the bypass. (**C**) Gene expression levels of the bypass chondrocyte markers. (**D**) Gene expression trends in the bypass. CPs, chondroprogenitors; BCs, bypass chondrocytes; HTCs, hypertrophic chondrocytes. (**E**) Representative differentially expressed genes between proliferative chondrocytes and bypass chondrocytes. (**F**) Gene expression levels of the pre-hypertrophic chondrocyte markers. (**G**) Immunostaining of the proliferative chondrocyte marker and the bypass chondrocyte marker. (**H**) Subclusters of the bypass chondrocytes.

Since both proliferative chondrocytes and bypass chondrocytes are able to differentiate into hypertrophic chondrocytes, these two subpopulations were further compared in details. The proliferative chondrocytes expressed higher levels of extracellular matrix genes such as *MATN3* and *COL6A3,* suggestive of their more mature characteristics. By contrast, the bypass chondrocytes exhibited higher *FGF2* and *GREM1* expressions therefore more resembling chondroprogenitors. The differential expression levels of *DKK1* and *WIF1* further confirmed that these two subpopulations were distinct from each other (Fig. 4E). However, the two subpopulations both expressed pre-hypertrophic chondrocyte markers, including *PTH1R, SMPD3* and *IHH,* demonstrating their hypertrophic destination (Fig. 4F). Interestingly, the immunostaining of the phalangeal epiphysis showed the presence of Neutral sphingomyelinase 2 (NSMase-2, product of the *SMPD3* gene) marked bypass chondrocytes in the SOC, where the type □ collagen (COL6) level was low, indicating that the typical endochondral ossification trajectory and the bypass may conduct chondrocyte hypertrophy in different ossification centers (Fig. 4G and S4B). In fact, distinctions were found between primary ossification center (POC) and SOC, including the timing, the location and the direction through which ossification proceeds(*28*). The chondroprogenitors and stem cell-like resting chondrocytes were located next to the secondary ossification center in mice, with no obvious proliferative zone in the middle(*29*), showing the different cell arrangement SOC from growth plate. Thus, the bypass indicated a direct ossification path in the SOC, which is distinguishable from the typical endochondral ossification in the growth plate, and explained why the typical proliferative zone can hardly be seen in the secondary ossification center.

The bypass chondrocytes were then further divided into two clusters (Fig. 4H). The left cluster (subcluster 1) which was close to the hypertrophic chondrocytes had higher expression levels of *SMPD3, RUNX2, MEF2C* and *PTH1R,* demonstrating the hypertrophic chondrocyte differentiation process, whereas the right cluster (subcluster 0) was enriched with *WIF1* and *CLU,* which were reported to regulate chondrocyte proliferation in osteoarthritis(*30*, *31*).

Taken together, these evidences demonstrated the direct ossification path that the chondroprogenitors directly differentiated to the hypertrophic chondrocytes in SOC, which could be quite different from canonical endochondral ossification process in growth plate.

In summary, we found that chondroprogenitors have two differentiation pathways to hypertrophic chondrocytes and one pathway to articular chondrocytes. Interestingly, as an alternative to the typical 4-stage endochondral ossification pathway, there was a direct ossification path by which chondroprogenitors could straightly differentiate into hypertrophic chondrocytes. Furthermore, our results revealed two new chondrocyte subpopulations (bypass chondrocytes in ossification path and *ID1^+^* chondroblasts in articular chondrocyte path) during postnatal epiphysis development in addition to six well-known subpopulations (chondroprogenitors, resting chondrocytes, proliferative chondrocytes, and hypertrophic chondrocytes in the endochondral ossification path; fibrochondrocytes, and superficial chondrocytes in the articular chondrocyte differentiation path). These results mapped a comprehensive developmental trajectory of chondrocyte differentiation in human epiphysis (Fig. 5), thereby expanding the knowledge of bone and articular cartilage, which could be utilized to design biotherapeutics for bone and articular cartilage regeneration.

**Figure 5.**
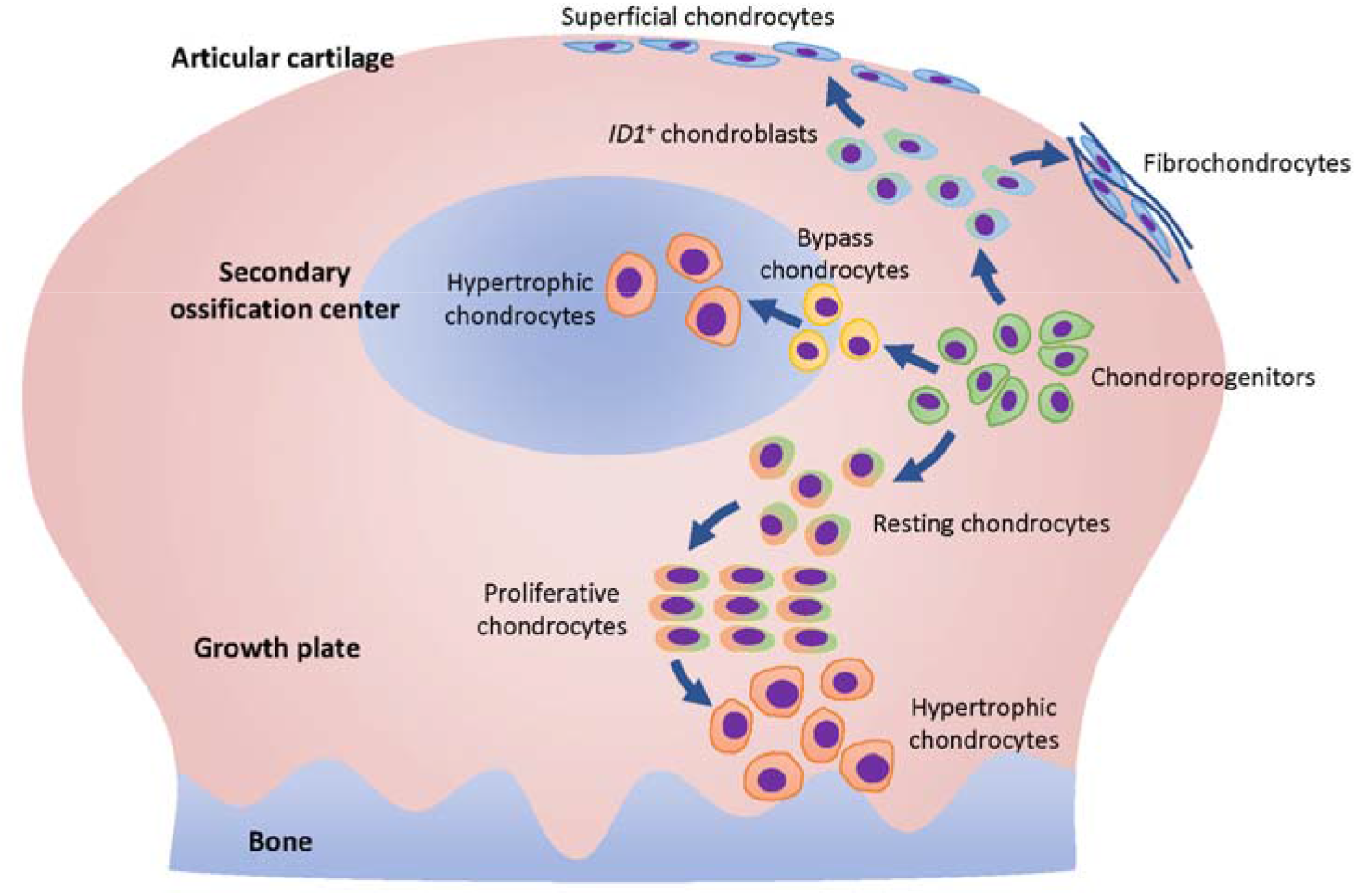
Schematic of the chondrocyte differentiation trajectories in human epiphysis.

## Data Availability

The sequencing raw data would be available before publication.

## Code Availability

The packages used for data analysis were stated in methods section. The full code for data analysis would be available before publication.

## Methods and materials

### 1. Human sample collection

We collected human polydactyl fingers from 4 babies (from 9 months to 8 years old). The procedure was approved by Children’s Hospital of Zhejiang University School of Medicine ethics committee (No. 2020-IRB-077). The bone and cartilage tissues were isolated and digested in 2% collagenases for 4 hours. The single cells were collected for 10× Genomics library preparation.

### 2 Preparation of single cell suspension

Cell number and viability were analyzed using hemocytometer and trypan blue. This method produces a single cell suspension with a concentration of 1000/μL and an activity exceeding 80%.

### 3. Single cell RNA sequencing: barcoding and cDNA synthesis

The single cell suspension was loaded onto a well on a 10x Chromium Single Cell instrument (10x Genomics). Barcoding and cDNA synthesis were performed according to the manufacturer’s instructions. Briefly, the 10x™ GemCode™ Technology partitions thousands of cells into nanoliter-scale Gel Beads-In-EMulsions (GEMs), where all the cDNA generated from an individual cell share a common 10x Barcode. Unique Molecular Identifier (UMI) was also added to identify the PCR duplicates. The GEMs were incubated with reverse transcription reagents to produce full length cDNA, which was then amplified via PCR to generate sufficient mass for library construction. The Qubit Fluorometer, Qubit dsDNA HS Assay Kit (Thermo Fisher Scientific, Cat# Q32854) and Agilent 2100 Bioanalyzer were used for QC and Qualitative analysis.

### 4. Single cell RNA sequencing: library construction and quality control

The cDNA libraries were constructed using the 10x Chromium™ Single cell 3’ Library Kit according to the manufacturer’s original protocol. Briefly, after the cDNA amplification, enzymatic fragmentation and size selection were performed using SPRI select reagent (Beckman Coulter, Cat# B23317) to optimize the cDNA size. P5, P7, a sample index and TruSeq read 2 (R2) primer sequence were added via End Repair, A-tailing, Adaptor Ligation, and PCR. The final single cell 3’ gene expression library contains a standard Illumina paired-end constructs (P5 and P7), Read 1 (R1) primer sequence, 16 bp 10x barcode, 12 bp UMI, cDNA fragments, R2 primer sequence and sample index. For post library construction QC and quantification, Qubit Fluorometer, Qubit dsDNA HS Assay Kit and Agilent 2100 Bioanalyzer were used.

### 5. Single cell RNA sequencing and generation of data matrix

Libraries were sequenced on an Illumina HiSeq X™ using HiSeq X™ Five Reagent Kit v2(Illumina, Cat# FC-502-2021).

### 6. Data processing

The .bcl files were called bases with Cellranger. The parameters were default. All the sequences in FASTQ were aligned to hg38.p5 reference and counted with Cellranger. The counts files were preprocessed with MATLAB. In detail, all fragments from non-protein coding genes were removed. All ribosome protein genes were removed because they have heavy multiple colinear effects interfering recognition of cell type with potential biological meaning. The counts data were normalized with CPM (counts per million), and all the cells with feature number lower than 1000 or higher than 7000 were removed. Cells with transcripts from mitochondrial genome occupying more than 50% in their library were removed. The genes whose CPM larger than two in at least 2 cells were selected for further analysis. The data were analyzed in R studio software (ver. 3.6.2), with Seurat package (ver. 3.1.2) from Satija Lab (https://satijalab.org/seurat/) and Monocle 3 package (ver. 0.2.2) from Cole Trapnell Lab (https://cole-trapnell-lab.github.io/monocle3/), following the standard protocol. In brief, the Seurat object was generated from digital gene expression matrices. Fourteen principal components were used in cell cluster with the resolution parameter set at 0.4. Then we performed cell cluster and UMAP Marker genes of each cell cluster were outputted to define cell clusters. The dimension reduction result conducted by Seurat was then used for pseudotime analysis by Monocle 3.

### 7. Tissue fixation and histology processing

Tissues for histology and immunostaining were fixed in 4% (w/v) paraformaldehyde for 24 h before decalcification in 10% (w/v) ethylene diamine tetraacetic acid (EDTA) solution. Subsequently, samples were embedded in paraffin and sliced (7 μm) for further safranin O/fast green staining or immunostaining.

### 8. Safranin O/fast green staining

The sections were deparaffinized and stained by hematoxylin for 20 min, followed by 8 min fast green staining, 1 s acetic acid washing, and 8 min safranin O staining subsequently. Then the slides were mounted by resinene and scanned with the digital scanner (3DHISTECH, Hungary)

### 9. Immunostaining

Paraffin sections for immunohistochemistry were treated with 0.25% trypsin (Gibco, USA), 3% (v/v) hydrogen peroxide in methanol, 1% (w/v) BSA, primary antibodies (TM4SF1, ab113504; follistatin, ab203131; COLX, ab58632; ID1, ab168256; COL1, AF7001; lubricin, ab28484; NSMase2, ab68735; COL6, ab6588) and secondary antibodies (A11008 and A21202, Invitrogen, USA) subsequently. The DAB substrate system (ZSGB-bio, China) was used for color development. Hematoxylin staining was utilized to reveal the cell nuclei. Then the slides were mounted with resinene and scanned by the digital slide scanner (3DHISTECH Pannoramic MIDI, Hungary). For immunofluorescence, the Alexa Fluor 488 or 546 conjugated secondary antibodies (Thermo Fisher Scientific, USA) were used, as well as DAPI (Beyotime, China) to reveal the cell nuclei. The images were acquired using a confocal microscope (Olympus, Japan).

## Acknowledgements

We give our special thanks to the patients who provided the precious samples for this research, as well as the families and doctors supporting them. We also would like to thank for the technical support by the Core Facilities, Zhejiang University School of Medicine.

## Author Contributions

H.S, Y.W and H.O designed the research; W.W and J.C collected the clinical samples; H.S, Y.W, T.Q, and Y.C performed the cellular and molecular experiments; H.S, T.Q, and C.A analyzed the data; H.S, T.Q, C.A, J.J, C.TG and H.O wrote the manuscript.

