## Supplemental Figures for "High-resolution architecture of human epiphysis formation"

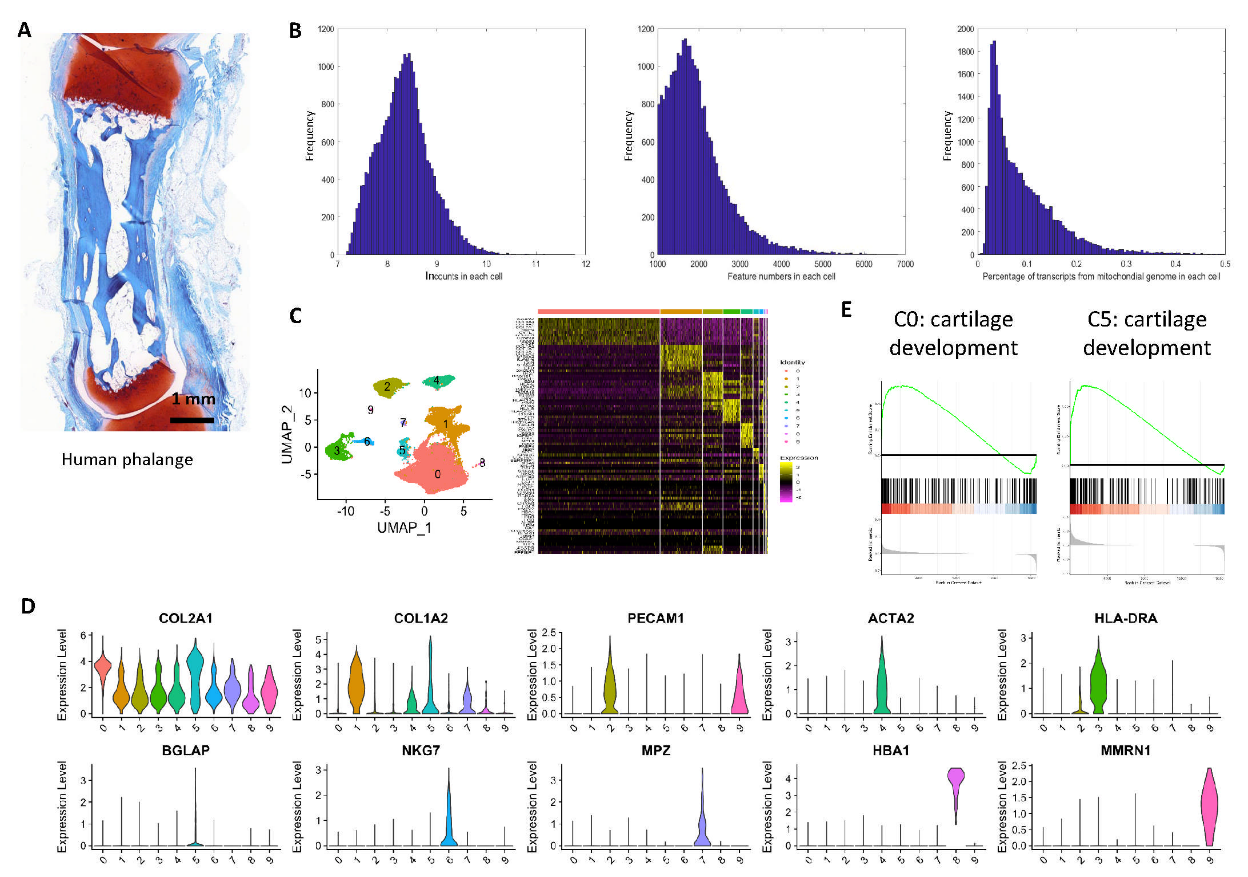

**Supplemental figure 1**: Human chondrocyte identification. (**A**) Safranin O staining of a 2-year-old human middle phalange. (**B**) Quality of the single-cell sequencing data. (**C**) Cell clustering result visualized by UMAP, and Heatmap of the differentially expressed genes for each cluster. (**D**) Marker genes of each cluster. (**E**) GSEA result for clusters 0 and 5.

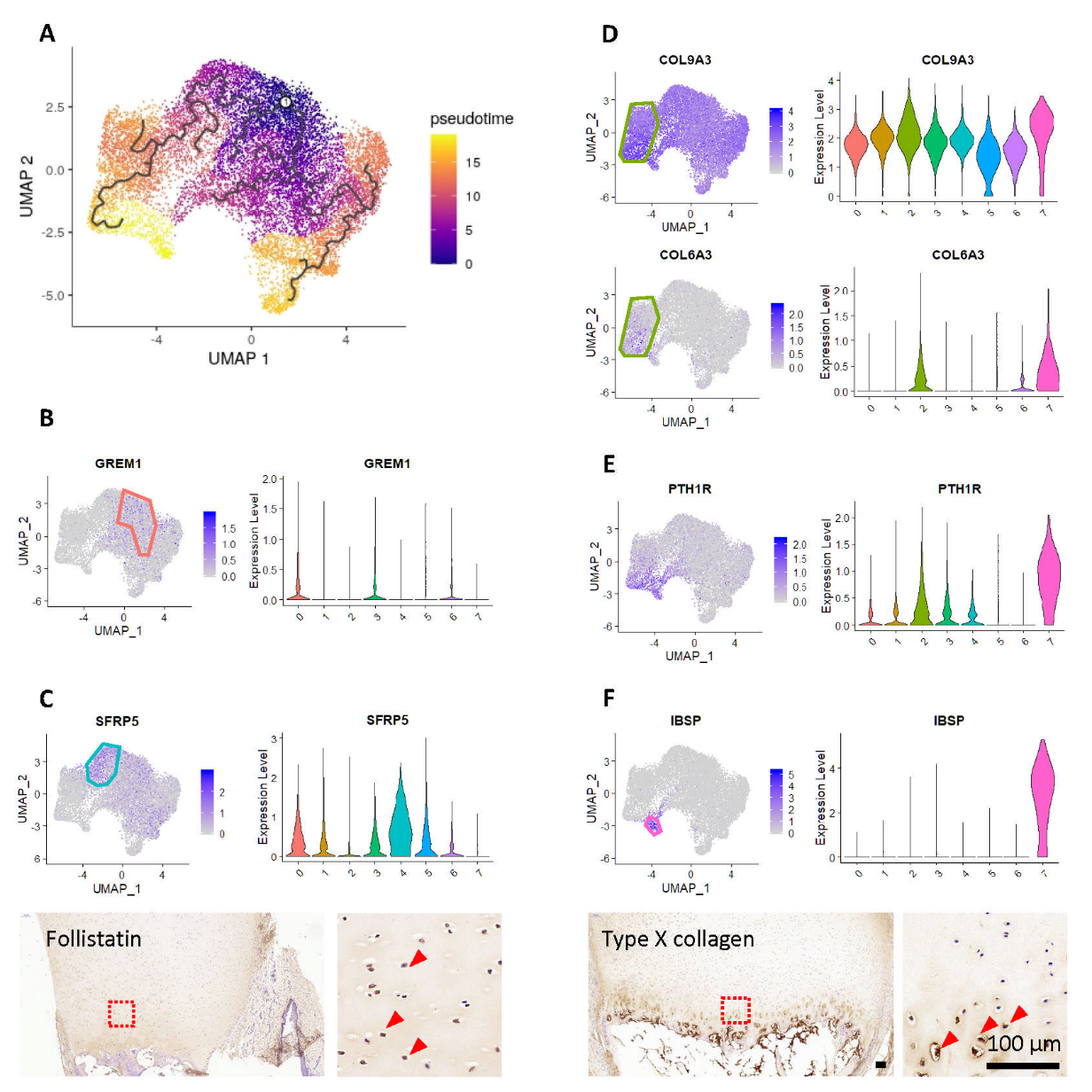

**Supplemental figure 2**: Identification of the typical endochondral ossification trajectory. (**A**) Pseudotime analysis visualized by UMAP. Red arrow shows the two directions of chondrocyte differentiation. (**B**) Gene expression levels of the chondroprogenitor marker. (**C**) Gene expression levels and immunostaining of the resting chondrocyte markers. (**D**) Gene expression levels of the proliferative chondrocyte markers. (**E**) Gene expression levels of the pre-hypertrophic chondrocyte marker. (**F**) Gene expression levels and immunostaining of the hypertrophic chondrocyte markers.

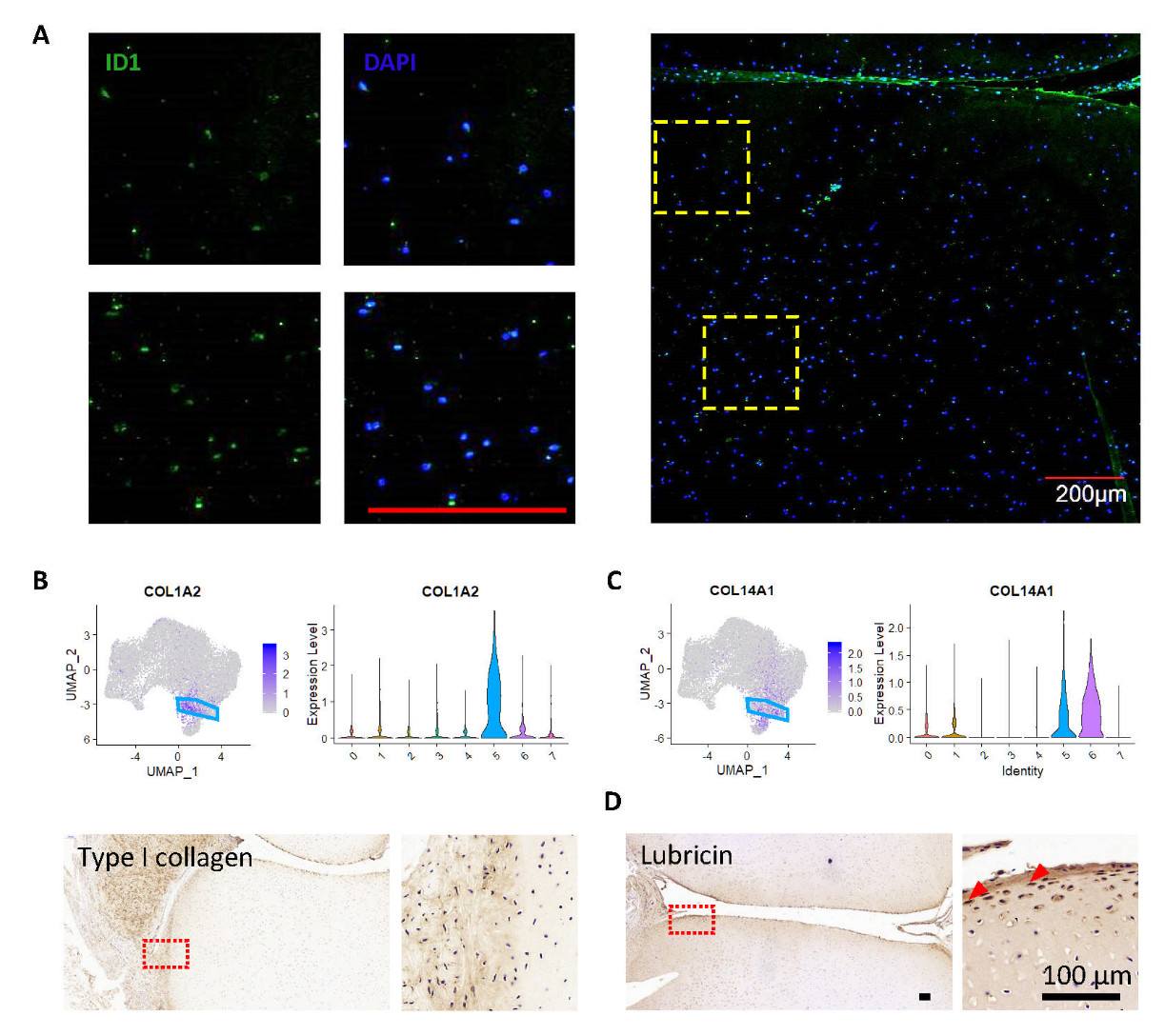

**Supplemental figure 3**: Identification of the articular chondrocyte differentiation trajectory. (**A**) ID1 immunostaining of the middle phalange. (**B** and **C**) Gene expression levels and immunostaining of the fibrochondrocyte markers. (**D**) Immunostaining of the superficial chondrocyte marker.

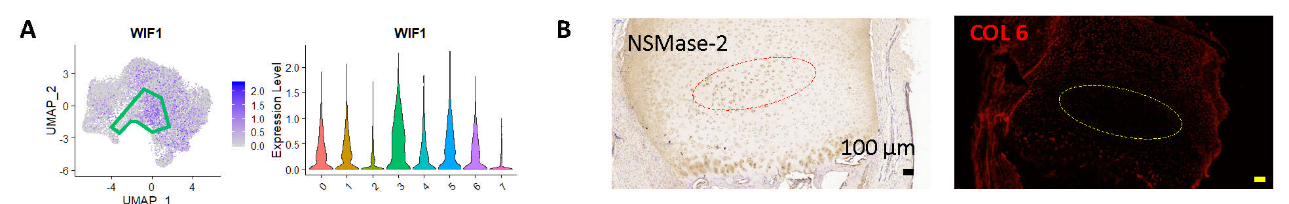

**Supplemental figure 4**: Identification of the chondrocyte differentiation bypass. (**A**) Gene expression levels of the bypass chondrocyte marker. (**B**) Immunostaining of the proliferative chondrocyte marker and the bypass chondrocyte marker. Low magnification.
